# Donor-Specific Digital Twin for Living Donor Liver Transplant Recovery

**DOI:** 10.1101/2025.02.21.639518

**Authors:** Suvankar Halder, Michael C. Lawrence, Giuliano Testa, Vipul Periwal

**Author notes:** Corresponding Author: Suvankar Halder.

## Abstract

Liver resection initiates a meticulously coordinated hyperplasia process characterized by regulated cell proliferation that drives liver regeneration. This process concludes with the complete restoration of liver mass, showcasing the precision and robustness of this homeostasis. The remarkable capacity of the liver to regenerate rapidly into a fully functional organ has been crucial to the success of living donor liver transplantation (LDLT). In healthy livers, hepatocytes typically remain in a quiescent state (G0). However, following partial hepatectomy, these cells transition to the G1 phase to re-enter the cell cycle. Surgical resection induces various stresses, including physical injury, altered blood flow, and increased metabolic demands. These all trigger the activation and suppression of numerous genes involved in tissue repair, regeneration, and functional recovery. Both coding and noncoding RNAs detectable in the bloodstream during this process provide valuable insights into the gene responses driving liver recovery.

This study integrates clinical gene expression data into a previously developed mathematical model of liver regeneration, which tracks transitions among quiescent, primed, and proliferating hepatocytes to construct virtual, patient-specific liver models. Using whole transcriptome RNA sequencing data from 12 healthy LDLT donors, collected at 14 time points over a year, we identified liver resection-specific gene expression patterns through Weighted Gene Co-expression Network Analysis (WGCNA). These patterns were organized into distinct clusters with unique transcriptional dynamics and mapped to model variables using deep learning techniques. Consequently, we developed a Personalized Progressive Mechanistic Digital Twin (PePMDT) for the livers of LDLT donors. The resulting PePMDT predicts individual patient recovery trajectories by leveraging blood-derived gene expression data to simulate regenerative responses. By transforming gene expression profiles into dynamic model variables, this approach bridges clinical data and mathematical modeling, providing a robust platform for personalized medicine. This study highlights the transformative potential of data-driven frameworks like PePMDT in advancing precision medicine and optimizing recovery outcomes for LDLT donors.

## 1 Introduction

The liver has the ability to regenerate and restore its functional mass following injury, disease, or surgical removal, such as partial hepatectomy. This regenerative capacity is vital for maintaining the body’s homeostasis, given its central role in numerous physiological processes, including detoxification of harmful substances, metabolism of nutrients and drugs, and the synthesis of essential proteins such as albumin and clotting factors [1–4]. The proliferation of mature hepatocytes, the functional cells of the liver, primarily drives liver regeneration. These hepatocytes, generally in a quiescent state (G0 phase of the cell cycle), can re-enter the cell cycle and proliferate in response to injury or loss of liver mass. This proliferation is a meticulously regulated process, ensuring that the liver regenerates to its original size without excessive or insufficient growth [5–7]. Apart from the hepatocytes, supporting endothelial cells, hepatic stellate cells, and Kupffer cells play critical roles in liver regeneration. Endothelial cells facilitate the restoration of the liver vascular network, ensuring adequate blood flow and nutrient delivery during the regenerative process. Hepatic stellate cells contribute by producing extracellular matrix components and growth factors, while Kupffer cells, the liver’s resident macrophages, secrete cytokines and chemokines that prime hepatocytes for regeneration [8–13]. This collective and coordinated interplay among hepatocytes and their supporting cells enables the capacity of the liver to regenerate while maintaining its essential functions. This capability of the liver to continue its normal functioning even during the regeneration process permits living donor liver transplantation (LDLT) [14].

Liver regeneration is influenced by multiple factors, including the host’s metabolic environment, graft type, and hepatectomy volume. Studies have shown that left lobe grafts regenerate faster than right lobes, with recipients exhibiting higher regeneration rates than donors [15]. Additionally, minor resections primarily rely on hypertrophy, whereas extensive resections involve both hypertrophy and hepatocyte proliferation, often leading to increased cell ploidy [15–17]. These complexities make it challenging to predict individual recovery trajectories following LDLT. Inter-patient variability significantly impacts recovery outcomes in LDLT, with factors such as genetic makeup, physiological status, pre-existing liver conditions, and immune responses playing crucial roles. For example, high intra-patient variability in tacrolimus levels has been associated with adverse outcomes, including graft rejection and the development of donor-specific antibodies [18]. Furthermore, the specialized nature of LDLT limits data availability, as it represents only a tiny fraction of total liver transplants. A study analyzing 73,681 adult recipients found that only 4% underwent LDLT, with substantial variation across transplant centers [19]. This scarcity of cases, combined with limited longitudinal follow-up data, poses significant challenges in developing clinically relevant predictive models for liver regeneration.

The process of liver regeneration is orchestrated by dynamic transitions of hepatocytes through quiescent, primed, and replicative states, regulated by intricate signaling pathways [20–22]. Understanding molecular drivers, including gene activation and suppression, is critical for modeling and predicting these regenerative dynamics [23]. Building on a foundation of experimental observations, we developed the first mathematical model to predict liver volume after partial hepatectomy (PHx) [24, 25], offering a quantitative framework to predict regenerative trajectories. To make such predictive models clinically relevant, we must integrate real-time patient-specific data with the dynamical framework provided by our dynamical model [26, 27]. This is essential for capturing the complex interplay of molecular, cellular, and systemic factors that drive liver regeneration [1, 21, 28].

However, challenges such as significant inter-patient variability, the nonlinear nature of liver recovery, and the integration of heterogeneous datasets complicate the development of personalized predictive models [27]. To address these complexities, this study introduces the Personalized Progressive Mechanistic Digital Twin (PePMDT), a donor-specific digital twin for the livers of LDLT donors. By integrating patient-specific clinical and molecular data with deep learning techniques, PePMDT enables precise simulation of regeneration dynamics. Similar deep learning ideas have also been applied in systems biology and multi-omics integration to uncover functional relationships between gene regulatory networks and physiological outcomes [29–32]. The digital twin framework begins with collecting blood samples from a specific LDLT donor after partial hepatectomy. Whole transcriptomic analysis is then performed, and the identified genes are clustered into various functional groups. These grouped genes are subsequently mapped onto our mechanistic mathematical model that describes liver regeneration processes. The model predicts dynamic variables at future time points, which are then used to forecast future gene expression patterns, enabling a data-driven approach to personalized liver recovery assessment, as depicted in Figure 1. This personalized framework has the potential to optimize post-transplant care, enhance recovery outcomes, and advance the field of regenerative medicine.

**Figure 1:**
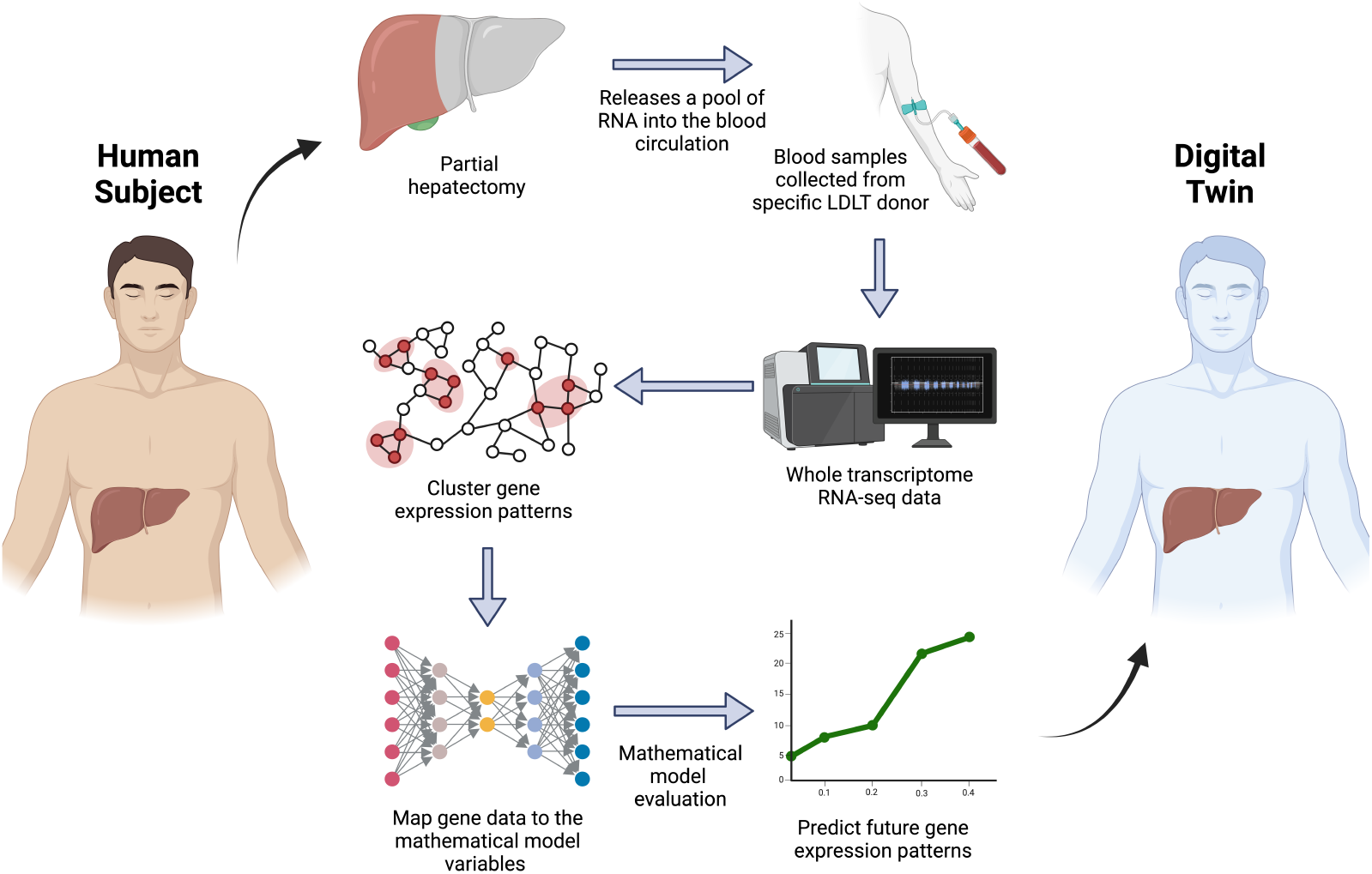
Framework of the digital twin. The general framework of the digital twin for liver regeneration. Blood samples are collected from a specific LDLT donor after partial hepatectomy, followed by whole transcriptomic analysis. Identified genes are clustered into functional groups and mapped onto a mechanistic mathematical model describing liver regeneration. The model predicts dynamic variables at future time points, which are then used to forecast future gene expression patterns, enabling a data-driven approach to personalized liver recovery assessment.

## 2 Data and Methods

### 2.1 Streamlining Gene Expression Profiles: Preprocessing Steps

Processed RNA-seq data were obtained from Lawrence *et al*. [33]. The dataset consists of whole-transcriptome RNA sequencing data from 12 healthy LDLT donors, collected at 14 predefined time points over a 1-year follow-up period. All LDLT donors underwent partial hepatectomy, and blood samples were collected at specific intervals post-surgery, including 5 minutes, 30 minutes, 1 hour, 2 hours, 3 hours, 4 hours, 1 day, 2 days, 3 days, 4 days, 10 days, 3 months, 6 months, and 12 months. The aim was to explore the dynamic transcriptional changes associated with liver regeneration post-PHx. This comprehensive dataset provided a unique opportunity to capture the liver’s gene expression landscape during recovery, from the immediate post-operative phase to the long-term restoration of liver function. Figure 2 presents a visual representation of the data structure, providing context for how the data was organized and utilized in this analysis. To ensure accurate downstream analysis and model predictions, the RNA-seq data underwent several preprocessing steps aimed at standardizing gene expression values across different patients and time points. The preprocessing steps were essential to handle potential technical and biological noise inherent in high-throughput sequencing data.

**Figure 2:**
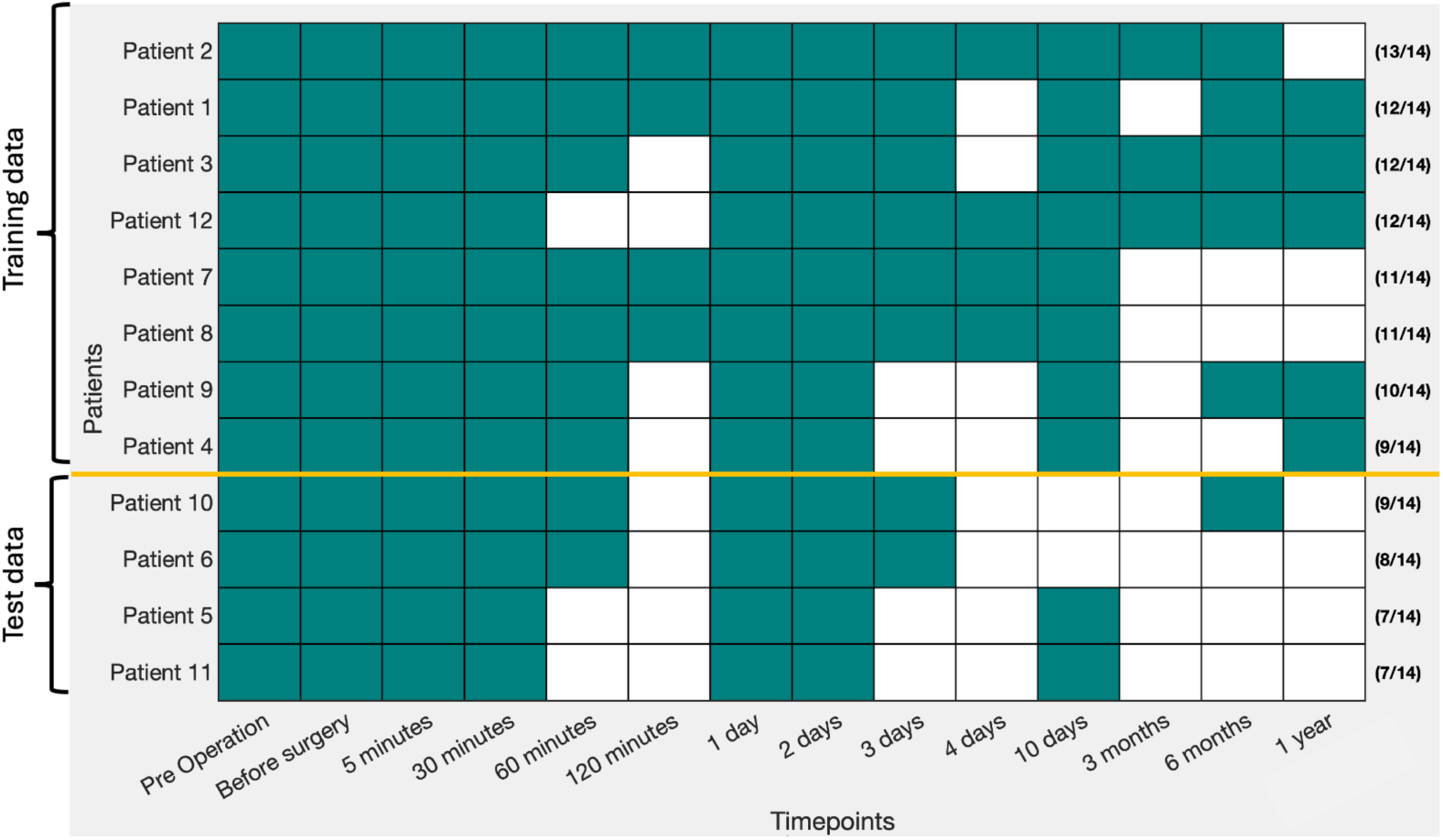
Data structure: The dataset consists of samples obtained from 12 healthy LDLT patients, collected at 14 distinct time points over the course of one year following partial hepatectomy. Filled color boxes indicate the presence of data, while blank spaces represent missing data. Patients are arranged in descending order based on the number of available time points. The number of available data points out of the 14 total time points is indicated on the right side for each patient. The top two-thirds of the dataset is used for training, while the remaining one-third serves as the testing dataset. The lower portion of the dataset was chosen for testing because these samples lack gene expression values in the later stages, providing an opportunity for predictive modeling.

### Missing Value Imputation

Due to the nature of RNA-seq data, some gene expression values were missing or below detection limits. To address this, we filled the missing values by averaging the corresponding expression values from other patients. This approach ensured that the imputed values were biologically plausible and aligned with the collective data structure. By using patient-averaged imputation, we preserved the continuity of expression profiles across time points and minimized biases in downstream analyses, ensuring robust and reliable integration of the data into our model.

### Correction of Negative Expression Values

In some instances, negative expression values were observed due to noise in low-expressed genes. These negative values were biologically unrealistic and could interfere with further analysis. To address this issue, we replaced all negative values with the average of the corresponding expression values from other patients. This replacement aims to ensure that no low or negative value artifacts skewed the results, particularly in downstream clustering or modeling.

### Log-Transformation

Gene expression values were log-transformed to reduce the skewness of the data and to stabilize the variance across time points and patients. Log-transformation compresses the range of gene expression values, potentially revealing more patterns in genes with both high and low expression. This transformation is especially useful in dynamic biological processes like liver regeneration, where gene expression changes can span several orders of magnitude over time. The transformation used was:

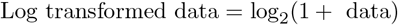

This preprocessing pipeline made the RNA-seq data ready for subsequent analyses, including clustering, temporal trend analysis, and mapping to the mathematical model variables.

### 2.2 Gene Co-expression Analysis: Identification of Temporal Gene Modules

The dataset contains 16,735 RNA gene transcripts in the blood, each expressed at least once during post-surgical recovery. Among these, 6,649 genes exhibited a differential expression of two-fold or more across at least two consecutive time points compared to the preoperative baseline following liver resection. Additionally, 1,110 genes showed expression changes of 1.5-fold or more at pre-resection time points, likely due to nonspecific surgical effects from the abdominal incision. These genes were excluded to focus on those potentially specific to liver regeneration. This left 5,876 genes that were identified as differentially expressed (DEGs) and specific to liver regeneration.

Weighted Gene Co-expression Network Analysis (WGCNA) [34] was employed on the DEGs to identify groups of genes that exhibited distinct temporal patterns during liver regeneration. These groups were further refined using self-organizing maps (SOM), which organized the genes into clusters based on their temporal expression profiles. SOM is particularly useful for visualizing time-series data, as it provides a low-dimensional representation of complex patterns, helping to reveal temporal trends in gene expression [35, 36]. These methods collectively identify clusters of highly correlated genes, which are believed to be involved in shared biological processes.

Each cluster is assigned a unique name and represents a set of genes whose expression patterns are highly similar across patients and time points. This method allows for the identification of clusters without prior knowledge of the biological pathways involved, making it an unbiased tool for analyzing dynamic gene expression data [34, 37].

### 2.3 Mathematical and Molecular Framework of Liver Regeneration

Liver regeneration following partial hepatectomy involves a series of biological processes, including hepatocyte proliferation, ECM degradation, and tissue remodeling. These processes are regulated by dynamic interactions between genes and the tissue environment, which can be modeled mathematically. Our previously developed mathematical model [24, 25] simulates liver regeneration by incorporating key processes such as liver volume recovery, hepatocyte proliferation rates, ECM turnover, and tissue remodeling.

The liver regeneration process is characterized by a coordinated transition of hepatocytes through distinct states—quiescent (Q), primed (P), and replicating (R). These transitions are illustrated in Figure 3a. While the total cell count consists of Q, P, and R cells, the overall liver volume increases due to the hypertrophy of primed and replicating cells [17]. The reduction in hepatocyte number following resection increases the metabolic load per hepatocyte (M/N), triggering the expression of growth factors (GF) and tumor necrosis factor (TNF). TNF promotes extracellular matrix (ECM) degradation via MMPs and induces IL-6 expression, which activates the JAK-STAT3 signaling pathway. STAT3, in turn, drives the transcription of SOCS3 and immediate early (IE) genes, shifting hepatocytes into a primed state. In this state, hepatocytes remain metabolically active but do not yet proliferate. Cytokines such as IL-6 and TNF-*α*, along with growth factors like HGF and EGF, further reinforce this priming by upregulating transcriptional programs involving immediate early genes, such as c-Jun and c-Fos. This prepares hepatocytes to respond to mitogenic signals and transition into the replicating state, where DNA synthesis and cell division occur to restore liver mass. As regeneration progresses, TNF levels decrease, ECM reformation occurs, and GF are inactivated, for example by uptake into the ECM, ultimately halting the primed-to-replicating transition and promoting the shift from replication back to quiescence. The molecular interactions underlying this process are depicted in Figure 3b. The detailed mathematical framework is given in [24, 25].

**Figure 3:**
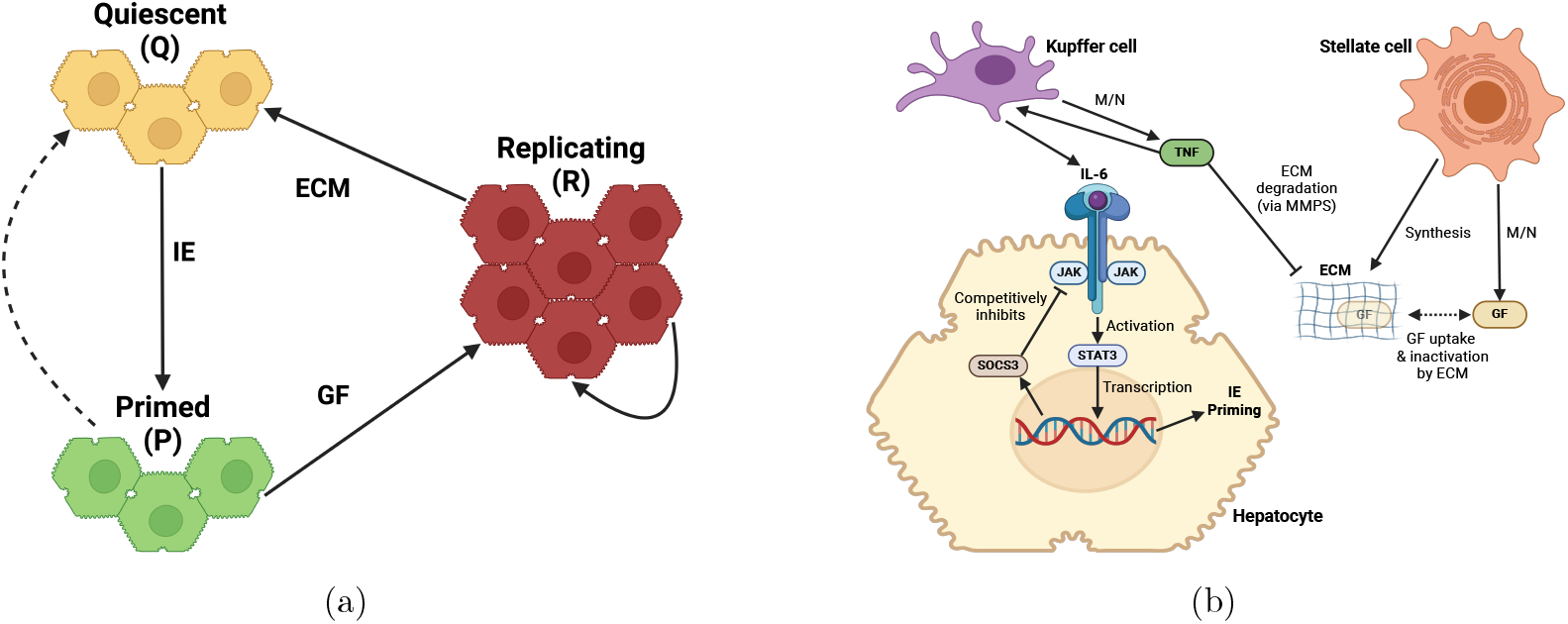
Schematic overview of liver regeneration. (a) The cell cycle dynamics during liver regeneration, highlighting changes in cell numbers. (b) The biochemical pathways involved in liver regeneration, illustrating key molecular interactions driving hepatocyte proliferation and tissue remodeling.

### 2.4 Constructing the Digital Twin

Integration of gene expression data with mechanistic modeling represents a potentially transformative approach to predicting personalized liver recovery. In this study, we constructed a digital twin, termed the Personalized Progressive Mechanistic Digital Twin (PePMDT), developed through a two-step mapping process combined with mathematical model simulations (see Figure 4). In the first step, we map gene expression data onto key physiological parameters of the mathematical model. This transformation enables us to represent biological states in terms of mathematically defined variables governing liver regeneration dynamics. Once the gene expression data has been mapped, we use the mathematical model to predict liver regeneration dynamics at future time points. These simulations predict the evolution of key physiological parameters, capturing hepatocyte proliferation, ECM turnover, and tissue remodeling over time. In the final step, we reverse-map the simulated mathematical model variables back to gene expression values at future time points. This process gives predicted gene expression profiles, allowing us to estimate the molecular responses of LDLT donors over time based on their initial gene expression states. These forward and reverse mappings are described in the following subsections.

**Figure 4:**
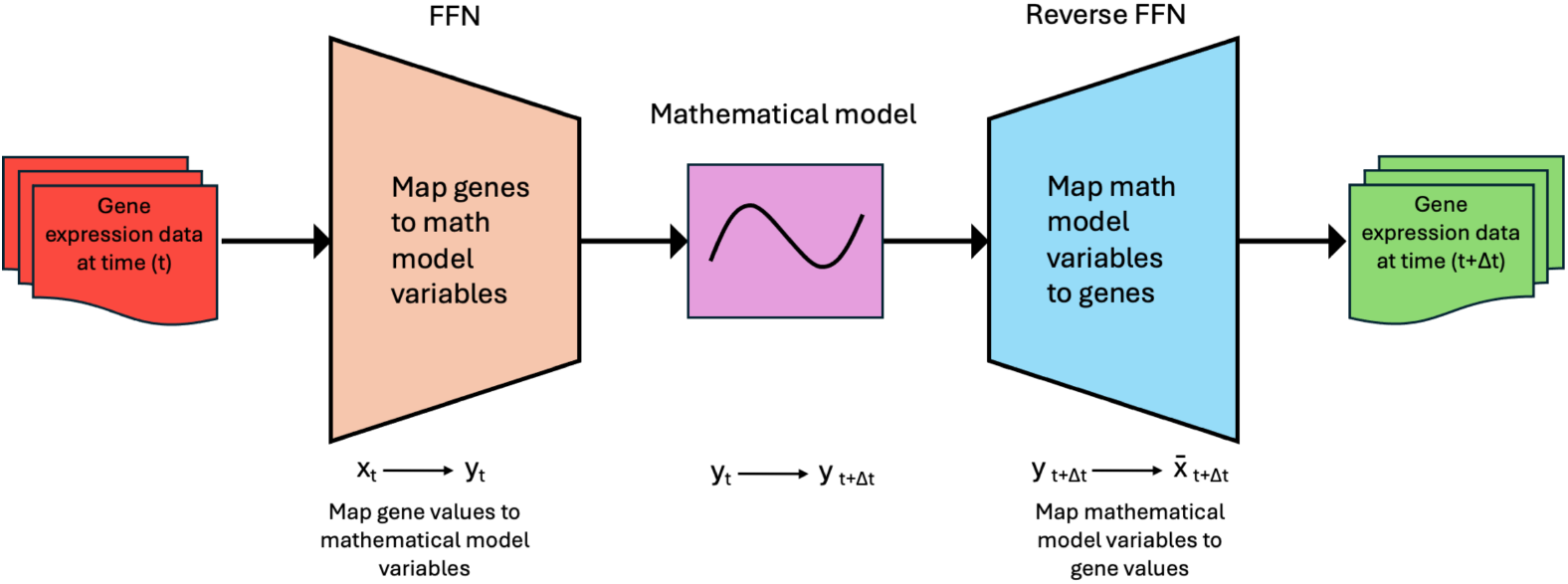
Schematic representation of the digital twin framework. The process involves a two-step mapping approach combined with mathematical model simulations. First, gene expression data is mapped onto key physiological parameters of the mathematical model. The model then simulates liver regeneration dynamics over future time points. Finally, the simulated mathematical values are reverse-mapped to reconstruct gene expression values, enabling the prediction of molecular responses in LDLT donors over time.

### Mapping Gene Expression Profiles to Mathematical Model Variables

The mathematical model [24, 25] describes hepatocyte state transitions in response to molecular signals, providing insights into liver recovery and potential therapeutic targets. However, they lack direct links to gene regulation. To bridge this gap, we used RNA-seq data from LDLT donors to map gene expression onto dynamic model variables. Using a feedforward neural network (FNN), we mapped gene expression data into key physiological parameters, enabling a gene-driven understanding of liver regeneration. The FNN consists of three fully connected layers, each followed by ReLU activation functions to introduce nonlinearity, potentially allowing the model to capture complex relationships between gene expression patterns and liver regeneration states. Dropout layers (0.2 probability) are incorporated to prevent overfitting, aiming to ensure robust generalization across different gene expression profiles. The input layer receives gene expression data clustered into biologically relevant groups (15 clusters), which is then transformed through progressively deeper layers (64 and 128 neurons) to extract meaningful patterns. The final layer outputs a lower-dimensional representation consisting of mathematical model variables corresponding to the measured gene expression levels computed based on patient-specific factors, such as the extent of liver resection, which plays a crucial role in the mathematical modeling framework. This parameter significantly influences the simulation of liver regeneration dynamics. The FNN is trained using a supervised learning approach, where the loss function (Mean Squared Error, MSE) measures the difference between predicted and actual physiological states. FNN parameters are optimized using backpropagation and the Adam optimizer [38].

### Reverse Mapping: Predicting Gene Expression from Model Variables

After mapping gene expression data to mathematical model variables, the model simulates liver regeneration dynamics over future time points. A reverse mapping step is required to translate these simulated variables back into gene expression values, predicting future molecular responses. This transformation is performed using a FNN as well. The architecture mirrors the FNN used for the forward mapping, consisting of three fully connected layers with ReLU activation functions to introduce nonlinearity and dropout layers (0.2 probability) to prevent overfitting. The input layer receives simulated model variables, processes them through progressively deeper layers (64 and 128 neurons), and outputs gene expression values corresponding to biologically relevant clusters. The model is optimized using the Adam optimizer, with MSE as the loss function to minimize the discrepancy between predicted and actual gene expression values.

## 3 Results

### 3.1 The Genetic Blueprint of Liver Regeneration: Clustering Analysis Reveals Temporal Gene Modules in Liver Regeneration

The combination of WGCNA and SOM [34–37, 39–42] was applied to identify temporally regulated gene modules, representing distinct biological processes across different phases of liver regeneration. This clustering approach allowed us to uncover gene networks with synchronized behavior, providing insights into the transcriptional programs driving recovery after partial hepatectomy.

After applying WGCNA and SOM (for details see Data and Methods section), the genes were clustered into 15 distinct groups (see Figure 5) - a higher number than the 8 clusters used by Lawrence et al. [33]. This increased number of clusters allowed for a more precise differentiation of DEGs based not only on their temporal expression patterns but also on their expression values, which proved beneficial for subsequent analyses. To assess the biological significance of the clusters, we compared our results with those of Lawrence *et al*. [33]. Consistent with their findings, we also identified that the clusters can be grouped into three prominent functional temporal gene modules corresponding to the main phases of liver regeneration: early response (Clusters 4, 5, 7, 8, 9, 10, 11, 12, 13, 14, 15), proliferation (Clusters 3, 6), and long-term recovery (Clusters 1, 2). Each module comprises genes with specific functions that are activated during distinct stages of the recovery process of the liver. SOM generates results based on random seeds, meaning that the outcomes may vary slightly for individual clusters with each run. However, this does not affect the overall functional modules of regeneration. The key processes associated with each module were determined using functional enrichment analysis using the Enrichr database [43] and are outlined to highlight their relevance to the overall biological context. The detailed figure of the Gene Ontology (GO) and pathway enrichment analysis are provided in the supplementary Figure 1.

**Figure 5:**
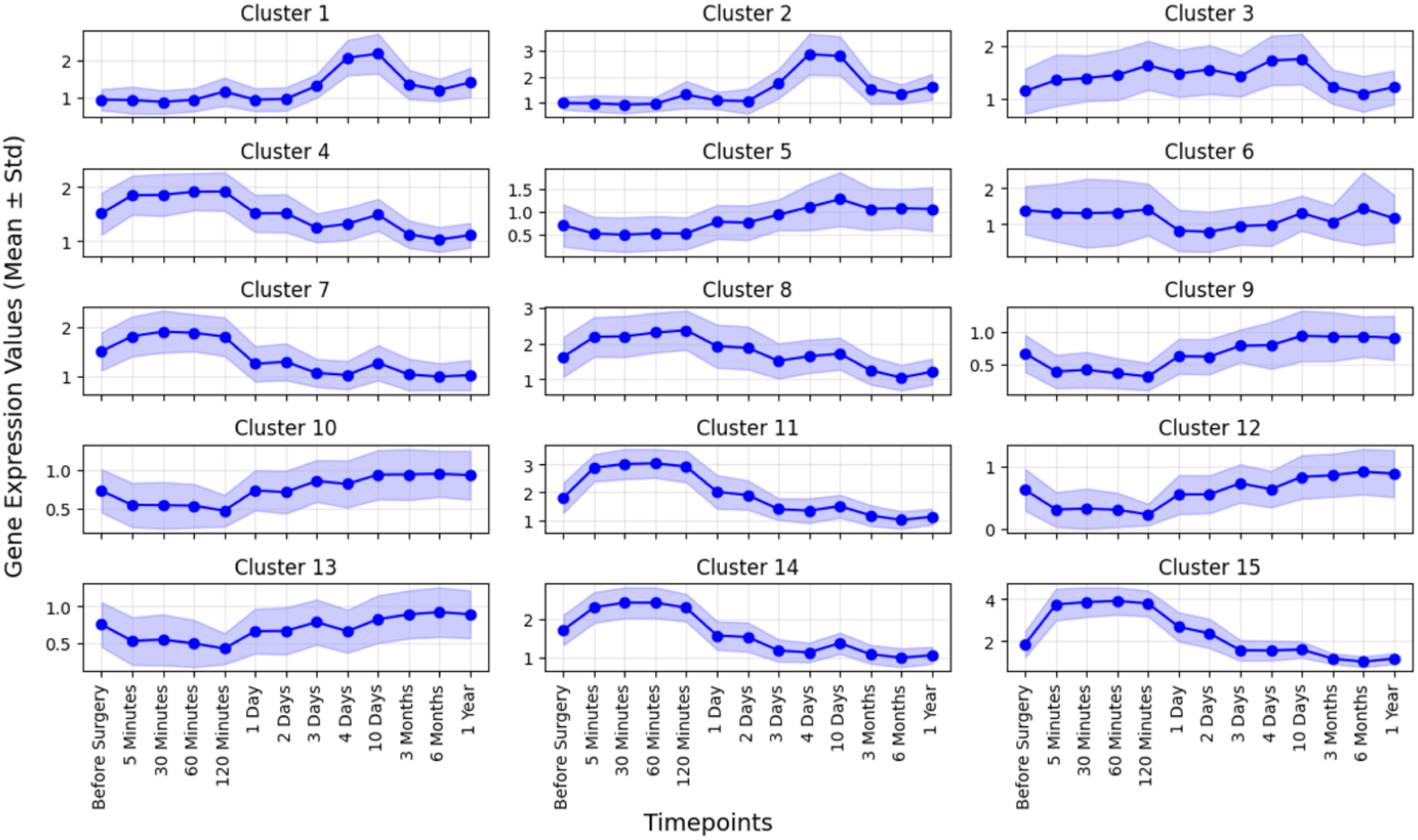
Temporal gene modules: A visual representation of the temporal gene expression patterns across 15 distinct clusters, which were subsequently grouped into 3 modules, during the liver resection and regeneration process. The bold lines represent the mean log2 fold change relative to preoperative values, and the shaded regions indicate the standard deviations.

#### Early-response module (Clusters 4, 5, 7, 8, 9, 10, 11, 12, 13, 14, 15)

This module consists of genes that peak within the first 24 hours post-PHx. These genes are primarily involved in the inflammatory response and early ECM degradation and remodeling. This triggers immune cell recruitment and matrix breakdown, which are crucial for initiating the regenerative process. Key genes in this module include those encoding cytokines, and chemokines responsible for ECM degradation. The temporal peak of this module aligns with the critical early stages of liver regeneration, where tissue damage needs to be repaired to create a favorable environment for cell proliferation.

#### Proliferation module (Cluster 3, 6)

Active between 1 and 10 days post-surgery, this module is enriched for genes related to cell cycle progression, hepatocyte proliferation, and DNA replication. During this phase, the liver undergoes rapid growth as hepatocytes proliferate to replace the lost tissue. This process is tightly regulated by growth factors and cell cycle regulators, ensuring a controlled proliferation. Genes in this module include cyclins, cyclin-dependent kinases (CDKs), and growth factor receptors. The peak of this module corresponds to the proliferative burst that occurs during liver regeneration, with hepatocyte replication reaching its highest levels between 3 and 10 days.

#### Long-term recovery module (Cluster 1, 2)

This module is characterized by genes that show sustained upregulation beyond 3 months post-PHx. These genes are associated with ECM organization, tissue remodeling, and the restoration of normal liver function. As the liver nears the completion of regeneration, it enters a phase of tissue stabilization, where the newly formed ECM becomes organized, and liver architecture is restored to support long-term functional recovery. The key genes in this module are involved in ECM remodeling, angiogenesis, and liver function maintenance, such as those encoding collagens, integrins, and enzymes involved in bile acid metabolism.

These three modules capture the temporal dynamics of liver regeneration and also reflect distinct biological processes that are activated sequentially.

### 3.2 Digital Twin: A Framework for Predicting Patient-Specific Future Gene Expression Patterns Related to Liver Mass Recovery with Precision

Leveraging the outcomes of both forward and reverse mapping, we developed PePMDT of liver regeneration, which simulates patient-specific recovery dynamics (see the Data and Methods section for details on its construction). This framework integrates gene expression data, mathematical model variables, and computational simulations, providing a comprehensive approach to predicting liver function and regeneration following PHx. Comparisons of predicted and actual gene expression profiles across multiple time points and patients demonstrated a strong alignment between the two datasets. In this analysis, the initial gene expression profiles of test patients (Before Surgery values) were input into the PePMDT model, which then predicted gene expression values for each cluster at subsequent time points. Figure 6 illustrates the results for all test patients: Patients 5, 6, 10, and 11.

**Figure 6:**
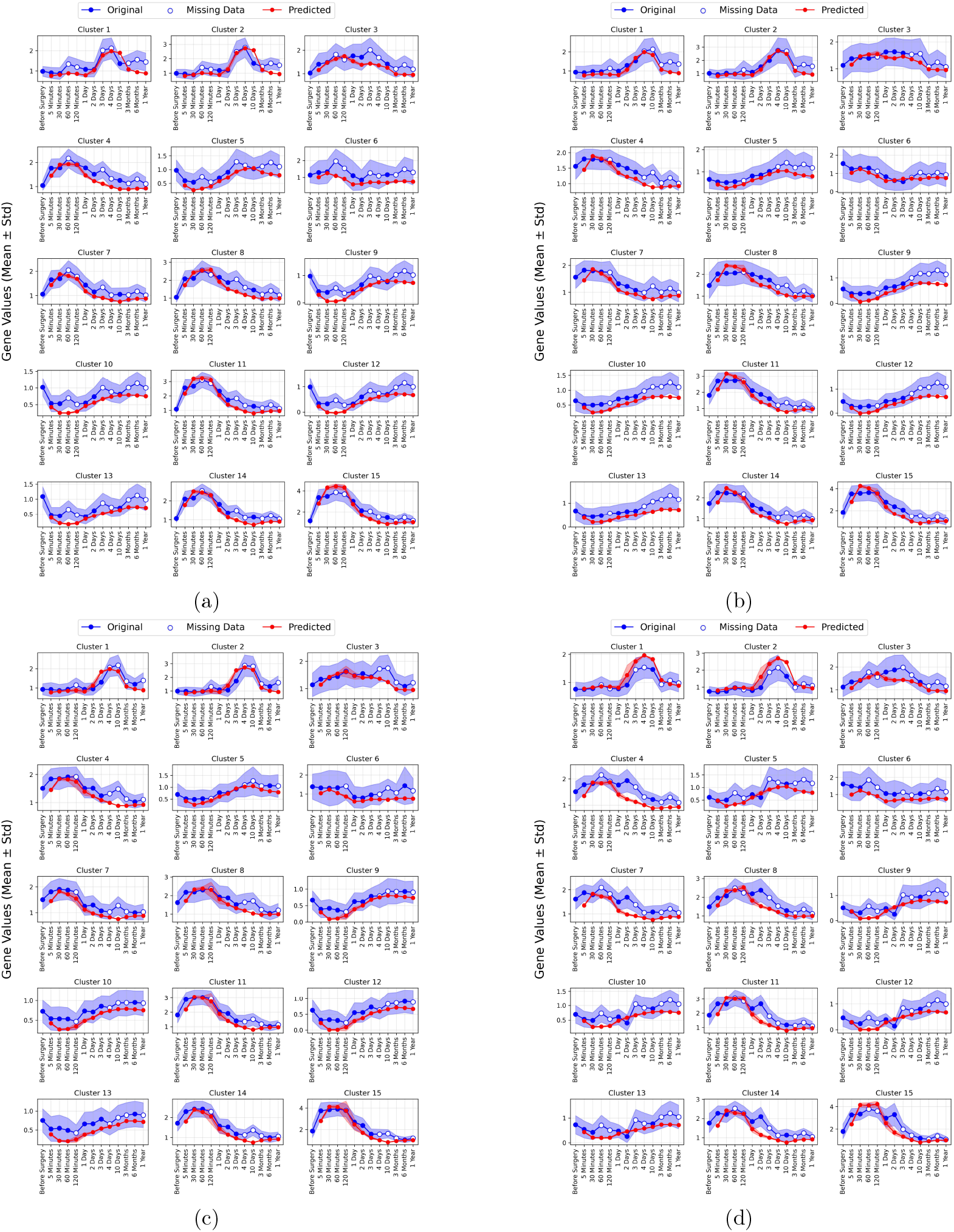
Comparison of original and predicted gene expression profiles. The plot illustrates the temporal gene expression patterns for Patient 5 (a), Patient 6 (b), Patient 10 (c), and Patient 11 (d). Bold lines represent the mean log2 fold change relative to preoperative values, with shaded regions indicating standard deviations. Red lines denote the predicted values, while blue lines represent the original (true) gene expression values. White dots indicate time points where data is missing in the original dataset.

To evaluate the accuracy of the predictions, we computed the cluster-wise MSE and Pearson correlation coefficient between the true and predicted gene expression profiles. For the majority of clusters, the MSE was found to be less than 0.3, with an average MSE across all test patients below 0.15. The Pearson correlation coefficient consistently exceeded 0.7, with the lowest average being 0.837, indicating a strong correlation between the predicted and true gene expression profiles. These results are depicted in Figure 7, where Panel A shows the cluster-wise MSE and Pearson correlation, and Panel B presents the averaged MSE and Pearson correlation across clusters.

**Figure 7:**
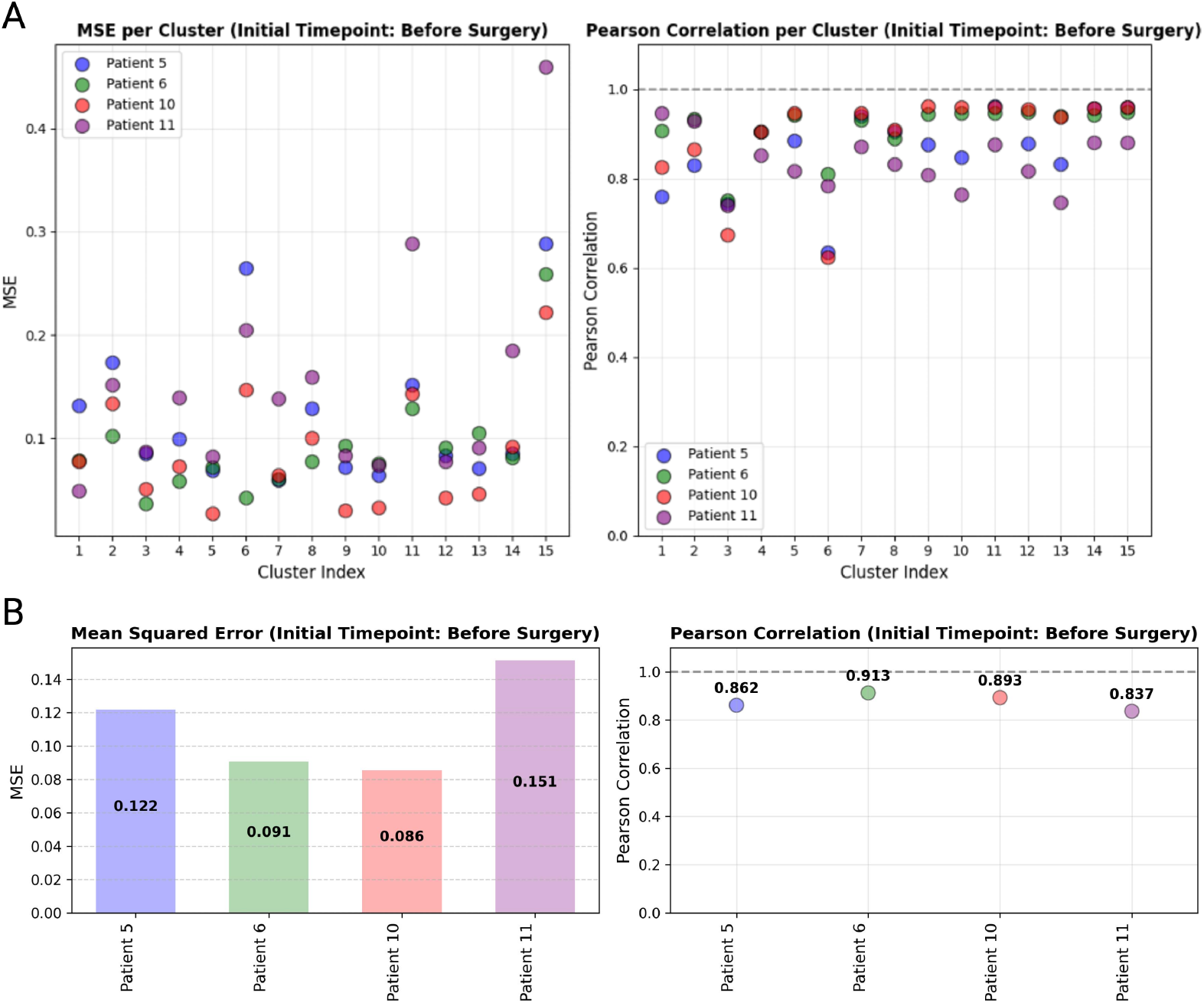
Patient-specific prediction accuracy. Comparison of prediction accuracy across individual test patients for the initial time-point (before surgery). Panel A shows the Mean Squared Error (MSE) and Pearson correlation coefficients between the true and predicted gene expression profiles for each cluster. Panel B displays the averaged MSE and averaged Pearson correlation across clusters.

To assess whether selecting a later time-point as the initial point improves prediction accuracy, we extended our analysis to two additional time-points: 5 minutes and 30 minutes post-surgery. However, we did not explore time-points beyond these, as the primary focus of our work is predicting future time-points based on earlier stages. We compared the results for these initial time-points and observed that the prediction accuracy remained consistent across all test patients. This comparison is shown in Figure 8, where the error bars represent the MSE between the true and predicted values across different time-points for each of the 15 clusters.

**Figure 8:**
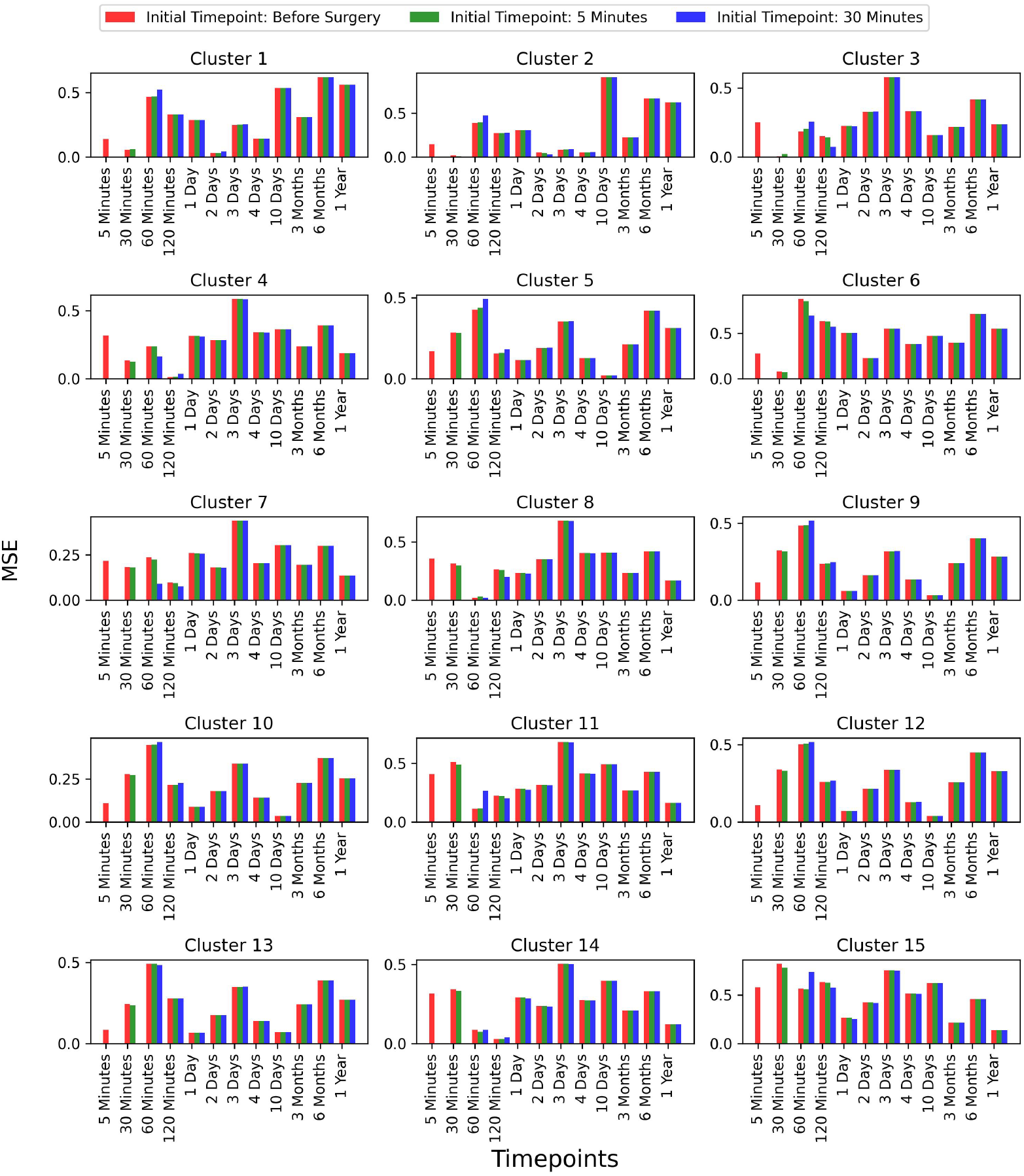
Comparison of prediction accuracy with different initial time-points (Continued figure: Part 1 for Patient 5). The figure depicts error bars representing the mean squared errors between the true values and the predicted values for each time-point for patient 5. The red, green, and blue bars correspond to the initial time-points chosen as before surgery, 5 minutes post-surgery, and 30 minutes post-surgery, respectively. This comparison is shown for all 15 clusters.

**Figure 8:**
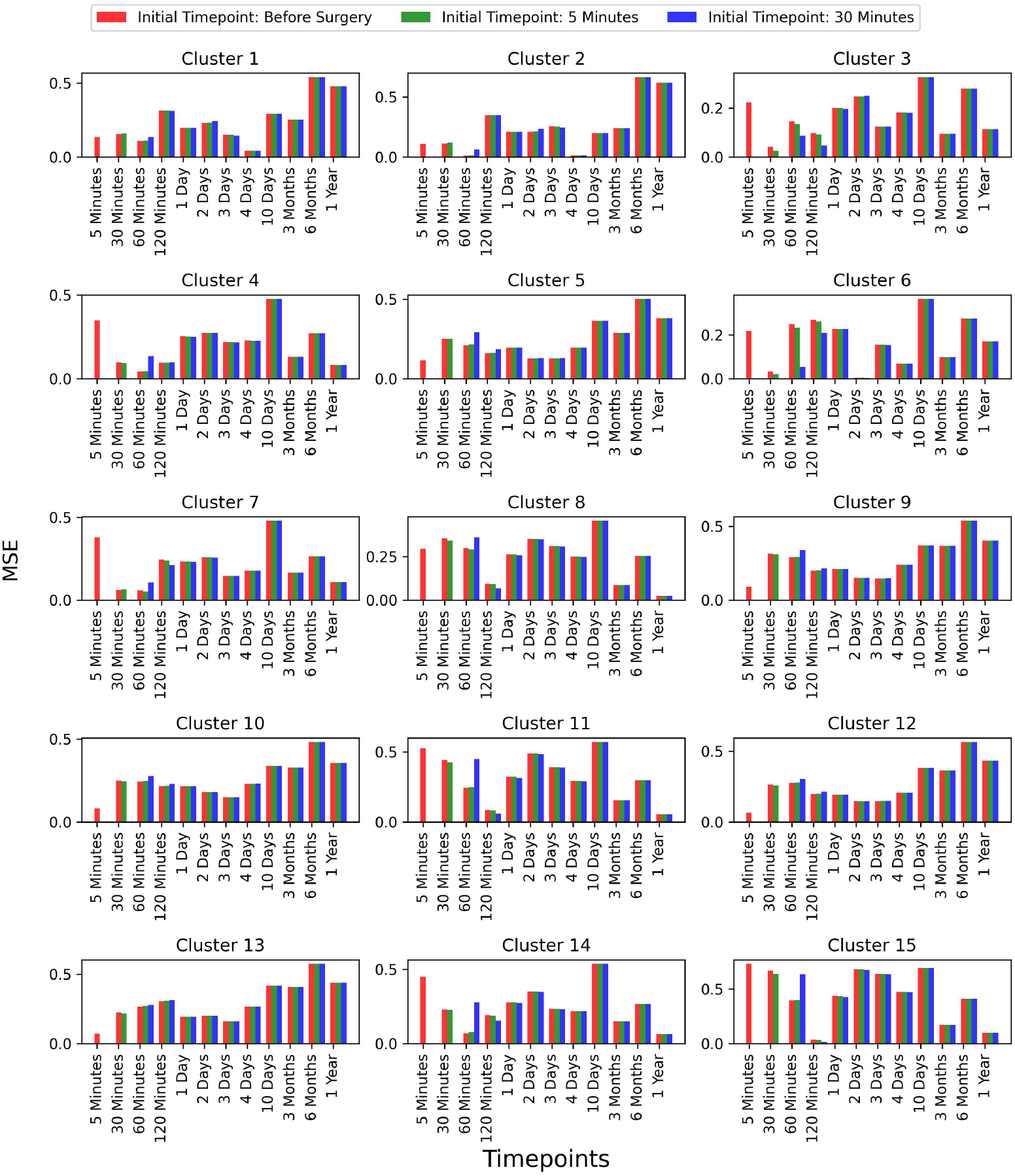
Comparison of prediction accuracy with different initial time-points (Continued figure: Part 2 for Patient 6). The figure depicts error bars representing the mean squared errors between the true values and the predicted values for each time-point for patient 6. The red, green, and blue bars correspond to the initial time-points chosen as before surgery, 5 minutes post-surgery, and 30 minutes post-surgery, respectively. This comparison is shown for all 15 clusters.

**Figure 8:**
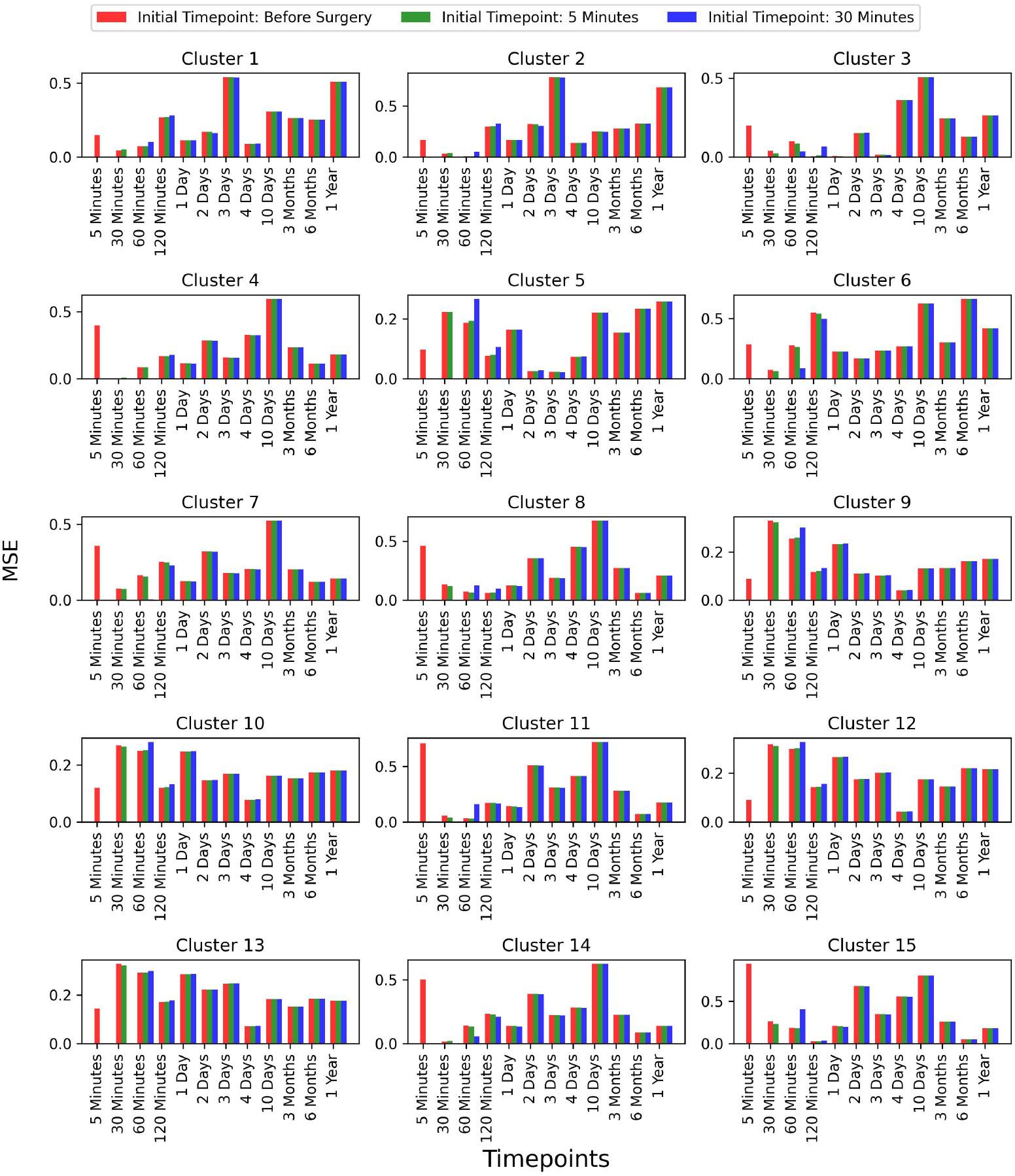
Comparison of prediction accuracy with different initial time-points (Continued figure: Part 3 for Patient 10). The figure depicts error bars representing the mean squared errors between the true values and the predicted values for each time-point for patient 10. The red, green, and blue bars correspond to the initial time-points chosen as before surgery, 5 minutes post-surgery, and 30 minutes post-surgery, respectively. This comparison is shown for all 15 clusters.

**Figure 8:**
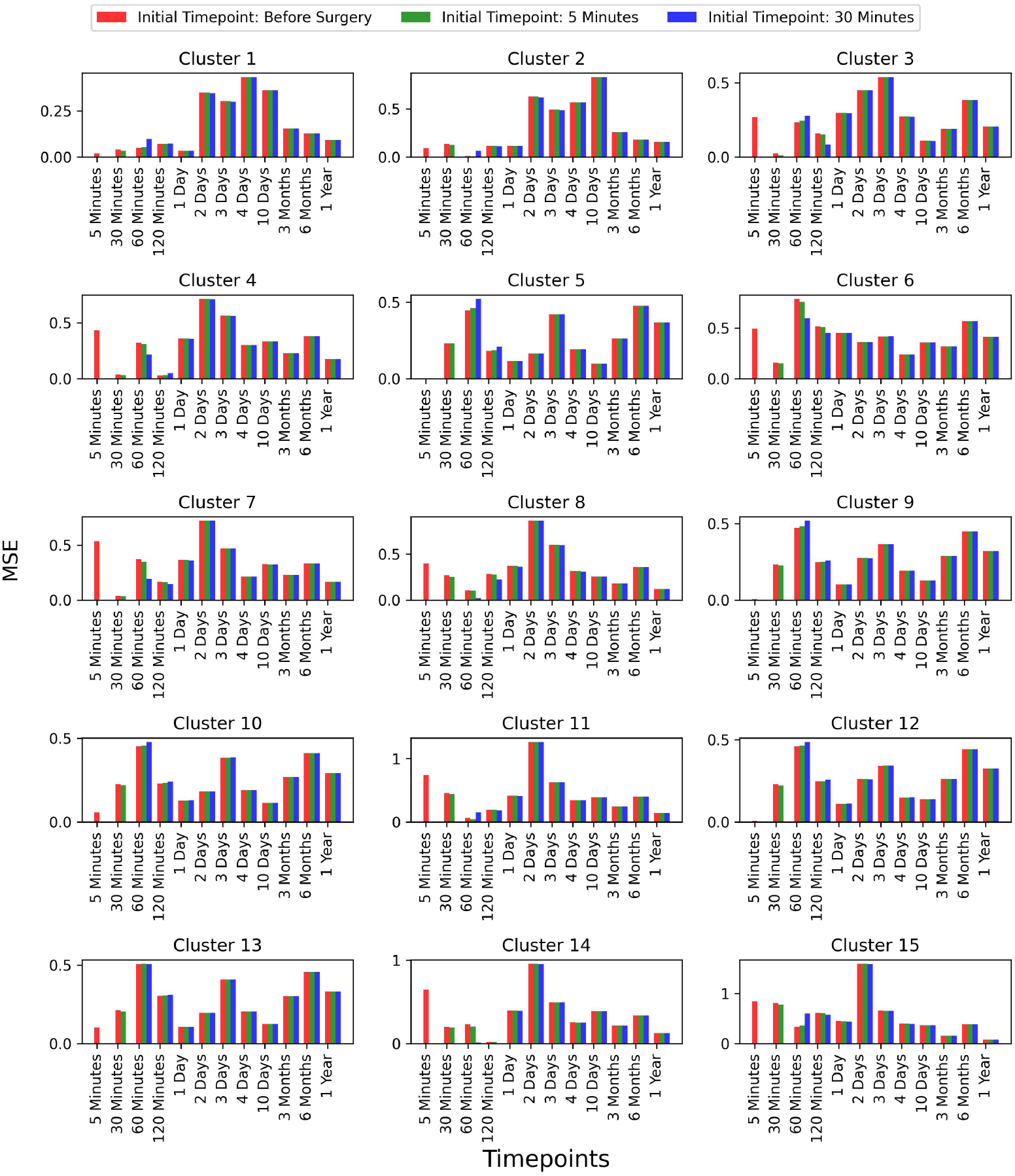
Comparison of prediction accuracy with different initial time-points (Continued figure: Part 4 (last part) for Patient 11). The figure depicts error bars representing the mean squared errors between the true values and the predicted values for each time-point for patient 11. The red, green, and blue bars correspond to the initial time-points chosen as before surgery, 5 minutes post-surgery, and 30 minutes post-surgery, respectively. This comparison is shown for all 15 clusters.

These results demonstrate that prediction accuracy does not significantly vary with different initial time-points. Although there is a small variation at the immediate next time-point for some clusters, no significant differences are observed at later time-points. This indicates that our model is capable of reliably predicting future gene expression values, even when the gene expression profile is provided as early as the before surgery time-point. To evaluate prediction accuracy for the post-surgical initial time-points (5 and 30 minutes), we computed the cluster-wise and average MSE and Pearson correlation coefficients between true and predicted gene expression profiles. The results, shown in Supplementary Figure 2, indicate strong correlations, reinforcing the model’s reliability in forecasting future gene expression patterns regardless of the initial time-point.

## 4 Discussion

Digital twin models have gained traction in healthcare, offering patient-specific simulations of disease progression and treatment responses [27]. Notable applications include heart failure management [44], personalized cancer treatment [45–49], organ transplantation [50], and liver regeneration [51, 52]. However, many existing models focus on static representations of disease progression, limiting their ability to capture the dynamic nature of biological processes. A notable exception is Ref. [53] which developed a framework for modeling personalized bladder cancer progression and optimizing adaptive multi-drug treatment strategies using deep reinforcement learning. However, no similar studies have been found for liver regeneration.

In this study, we developed PePMDT, a digital twin model that integrates deep learning with mathematical modeling to predict patient-specific gene expression dynamics during liver regeneration. Our approach successfully reconstructed future postoperative gene expression profiles using very early postoperative data, including pre-resection data as well as data from 5 minutes and 30 minutes after resection, demonstrating high predictive accuracy. PePMDT offers a dynamic, personalized framework that seamlessly integrates gene expression data with mechanistic simulations to predict long-term liver function recovery.

To assess the predictive accuracy of PePMDT, we evaluated its performance on patient-specific data. Each patient exhibited unique gene expression patterns, underscoring the personalized nature of liver regeneration. Our model accurately predicted gene expression profiles across all patients, while also capturing substantial variability in recovery trajectories. Quantitatively, we assessed prediction accuracy using mean squared error and Pearson correlation coefficients across gene clusters, with the lowest average correlation across all patients being 0.83, indicating strong alignment between observed and predicted values. Beyond its predictive capability, PePMDT enables what-if simulations, allowing the exploration of liver recovery under different clinical scenarios, such as varying degrees of liver damage or resections. This predictive power provides a valuable tool for understanding liver regeneration at a molecular level and optimizing treatment strategies. By constructing a computational model informed by real patient gene expression data, PePMDT facilitates personalized medicine applications, allowing clinicians to anticipate complications and tailor interventions based on a patient’s molecular profile. Moreover, simulations of gene expression dynamics over time validate the model’s utility in predicting individual recovery trajectories. By accurately forecasting future gene expression states based on early molecular patterns, PePMDT bridges high-throughput genomic data with physiologically relevant mathematical modeling, offering a scalable and robust framework for precision medicine in liver regeneration.

## Acknowledgments

This research was supported by the Intramural Research Program of the NIH, the National Institute of Diabetes and Digestive and Kidney Diseases (NIDDK, Project Z01-DK075059).

## Notes

### Competing Interest Statement

The authors have declared no competing interest.

