## Supplementary materials for "Donor-Specific Digital Twin for Living Donor Liver Transplant Recovery"

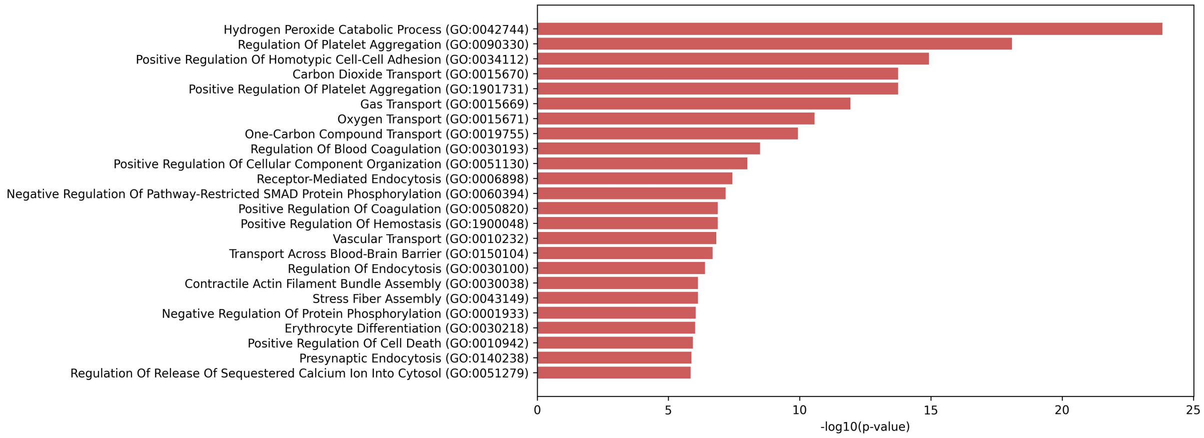


(a) Cluster 1

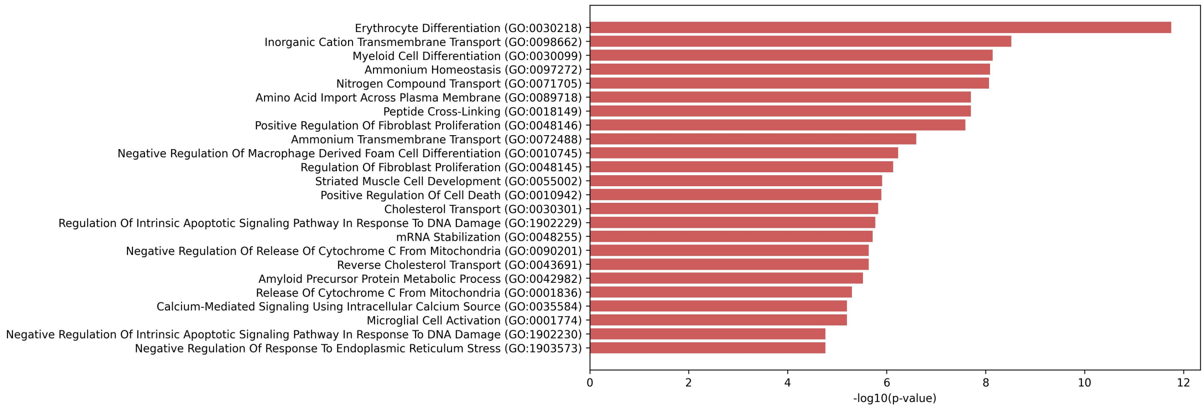


(b) Cluster 2

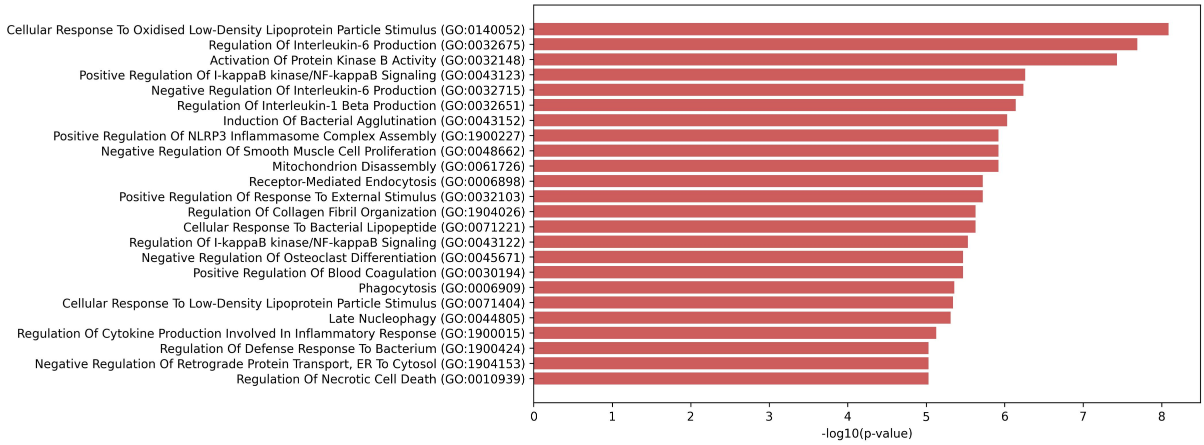


(c) Cluster 3

Figure 1: **Gene Ontology (GO) and pathway enrichment analysis of differentially expressed genes (DEGs) (Continued figure: Part 1).** The most significantly enriched GO terms identified using Enrichr analysis are shown. GO terms with p-values $< 10^{-5}$ were considered. The x-axis represents -log10(p-value), indicating the significance of enrichment. GO terms related to biological processes, molecular functions, and cellular components provide insights into key pathways influenced by DEGs.

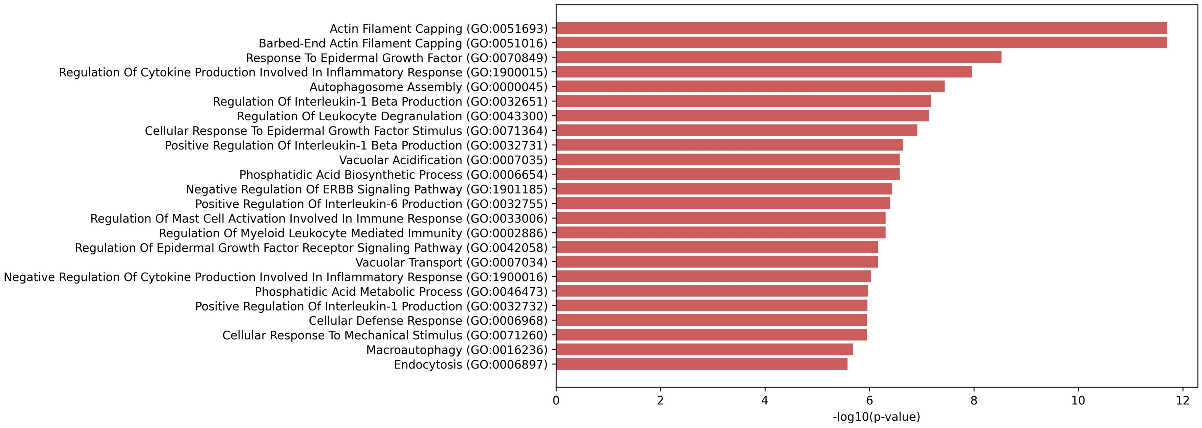


(d) Cluster 4

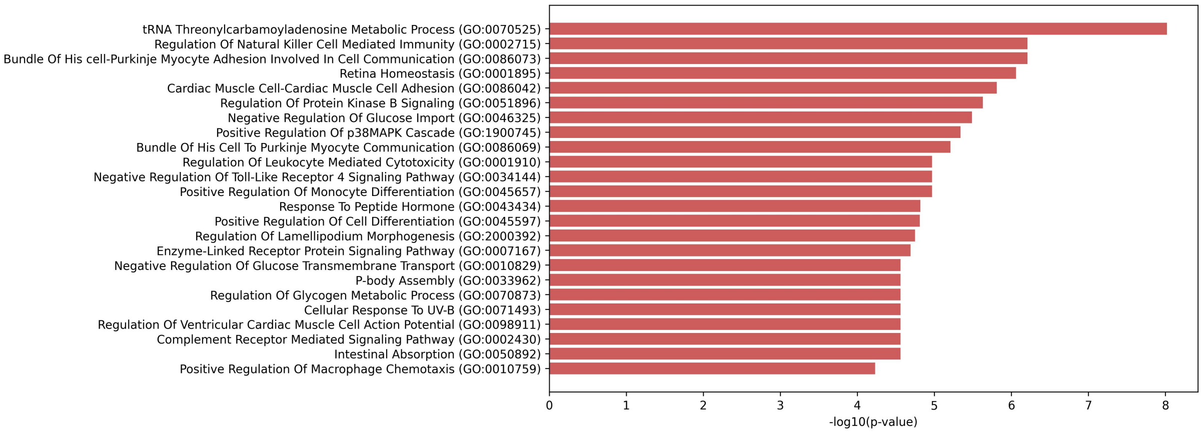


(e) Cluster 5

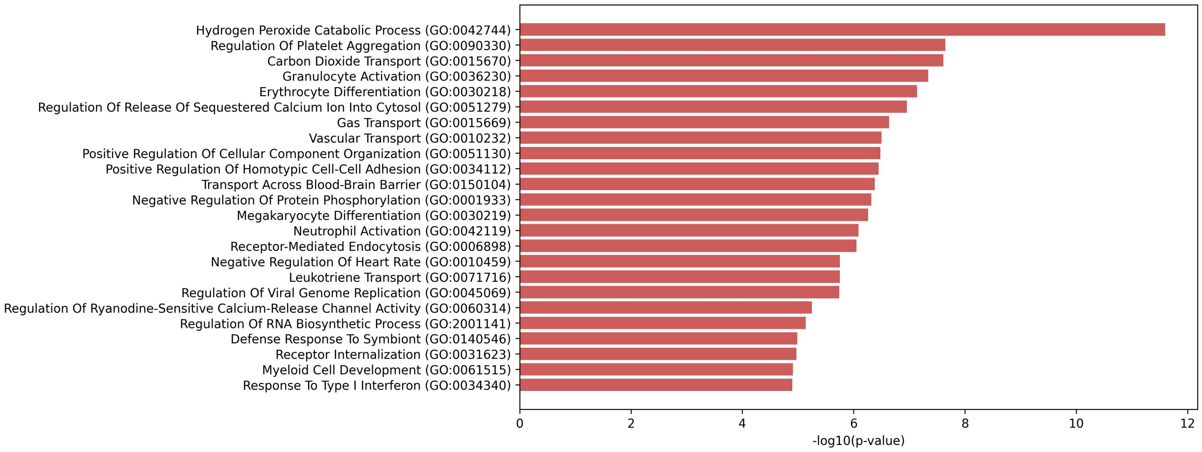


(f) Cluster 6

Figure 1: **Gene Ontology (GO) and pathway enrichment analysis of differentially expressed genes (DEGs) (Continued figure: Part 2).** The most significantly enriched GO terms identified using Enrichr analysis are shown. GO terms with p-values $< 10^{-5}$ were considered. The x-axis represents -log10(p-value), indicating the significance of enrichment. GO terms related to biological processes, molecular functions, and cellular components provide insights into key pathways influenced by DEGs.

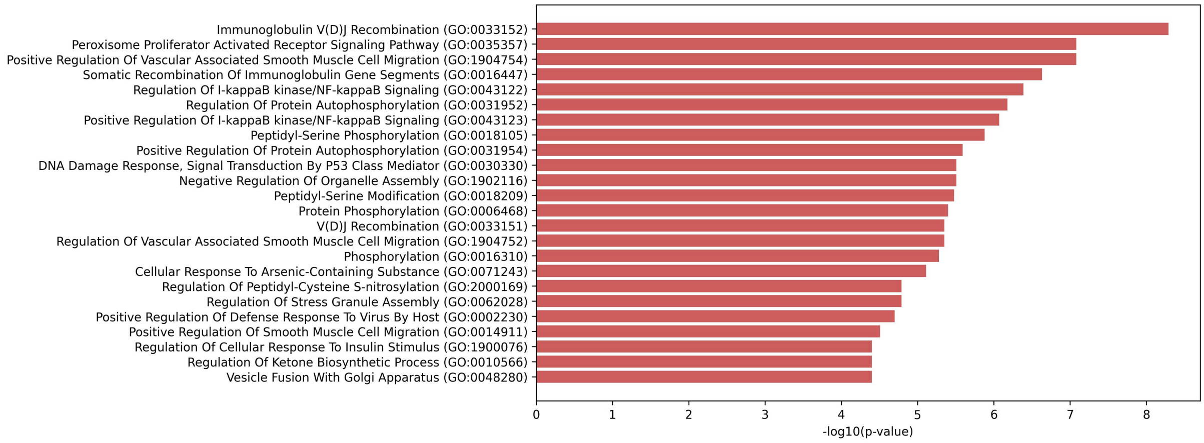


(g) Cluster 7

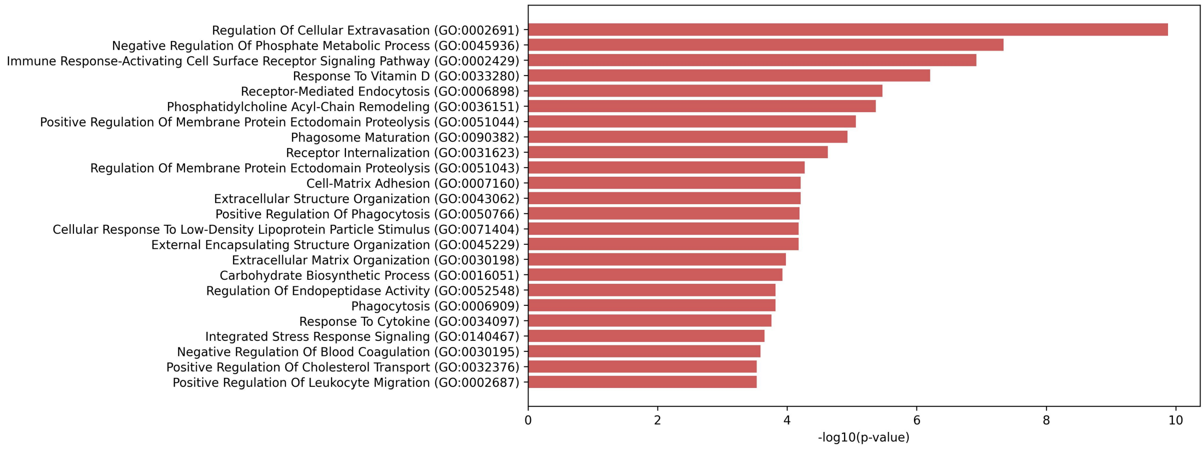


(h) Cluster 8

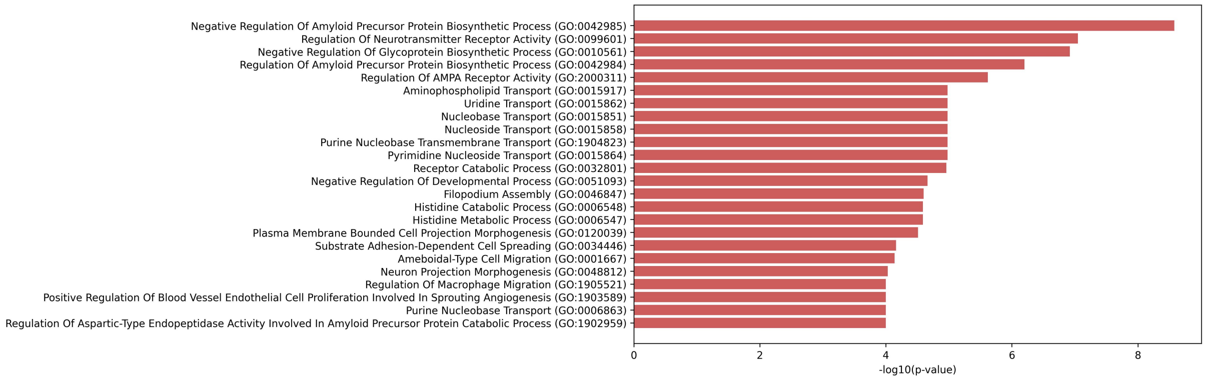


(i) Cluster 9

Figure 1: **Gene Ontology (GO) and pathway enrichment analysis of differentially expressed genes (DEGs) (Continued figure: Part 3).** The most significantly enriched GO terms identified using Enrichr analysis are shown. GO terms with p-values $< 10^{-5}$ were considered. The x-axis represents -log10(p-value), indicating the significance of enrichment. GO terms related to biological processes, molecular functions, and cellular components provide insights into key pathways influenced by DEGs.

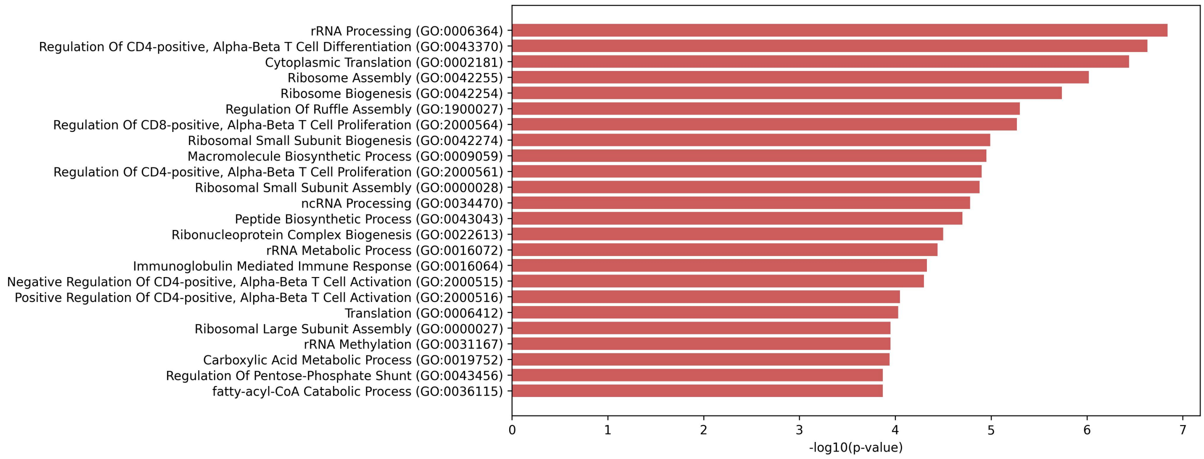


(j) Cluster 10

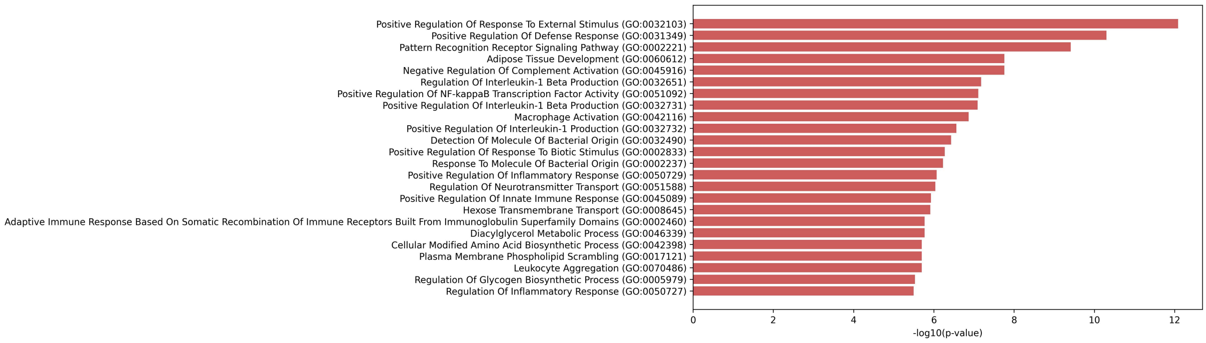


(k) Cluster 11

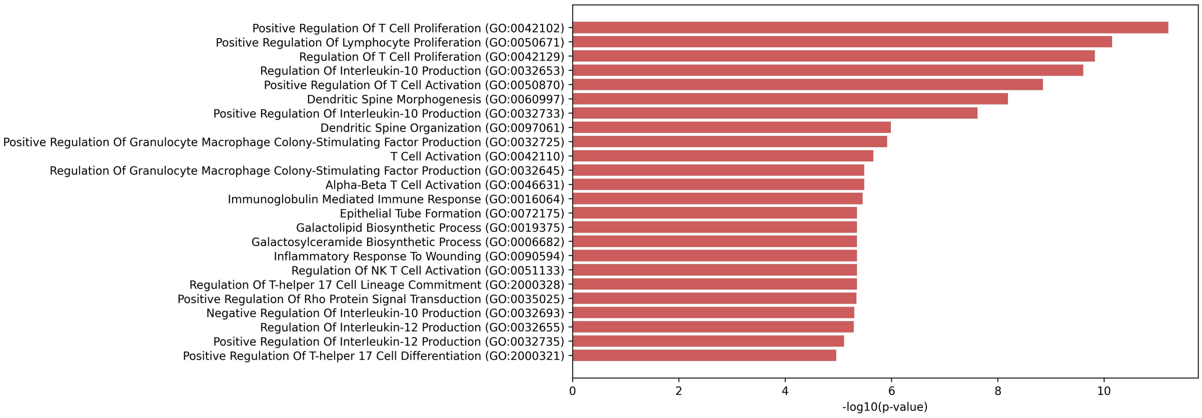


(l) Cluster 12

Figure 1: **Gene Ontology (GO) and pathway enrichment analysis of differentially expressed genes (DEGs) (Continued figure: Part 4).** The most significantly enriched GO terms identified using Enrichr analysis are shown. GO terms with p-values $< 10^{-5}$ were considered. The x-axis represents -log10(p-value), indicating the significance of enrichment. GO terms related to biological processes, molecular functions, and cellular components provide insights into key pathways influenced by DEGs.

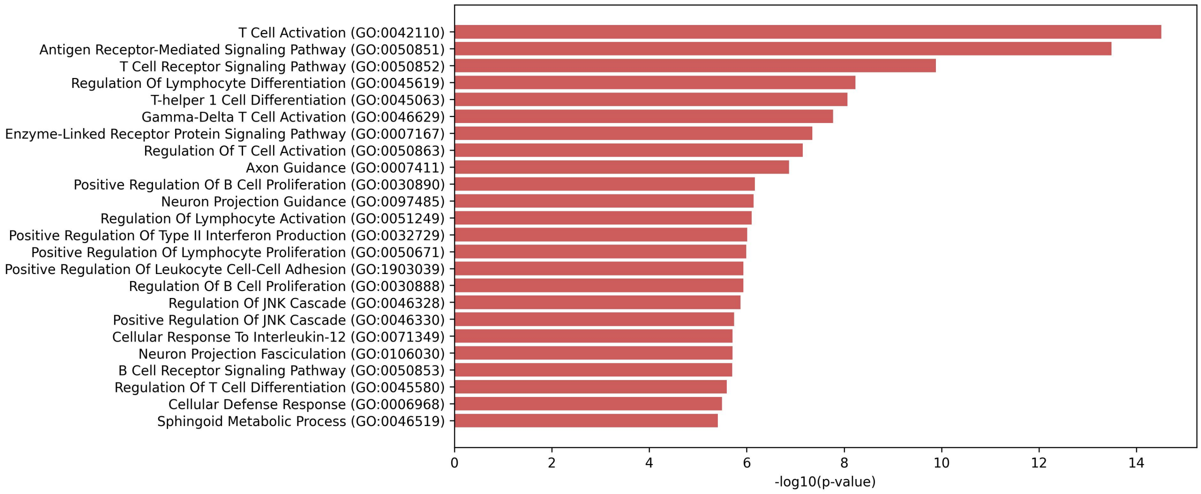


(m) Cluster 13

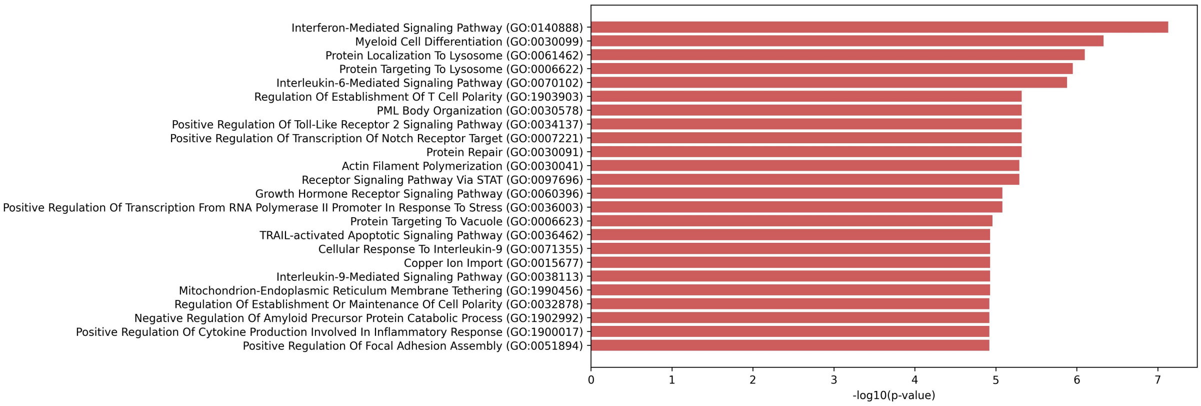


(n) Cluster 14

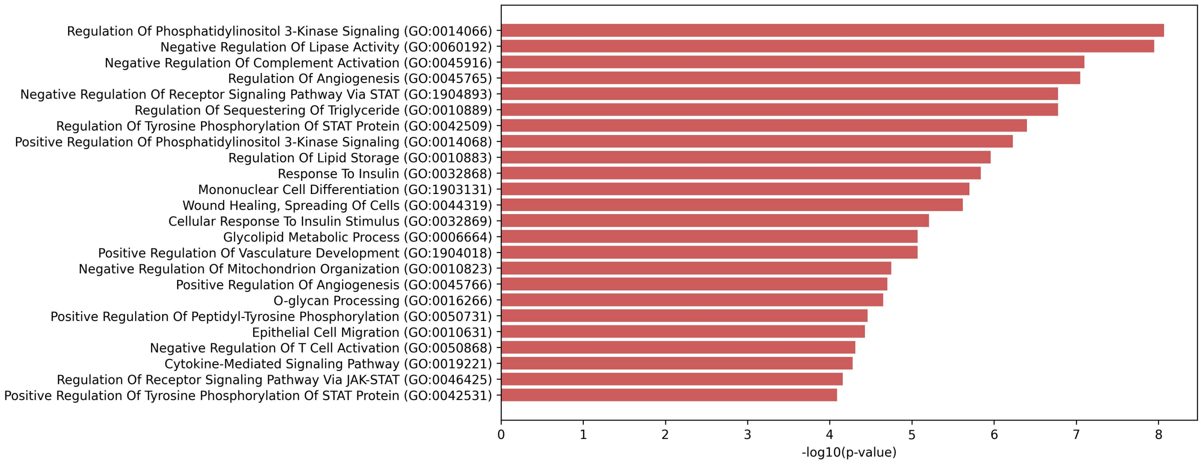


(o) Cluster 15

Figure 1: **Gene Ontology (GO) and pathway enrichment analysis of differentially expressed genes (DEGs) (Continued figure: Part 5 (last)).** The most significantly enriched GO terms identified using Enrichr analysis are shown. GO terms with p-values $< 10^{-5}$ were considered. The x-axis represents -log10(p-value), indicating the significance of enrichment. GO terms related to biological processes, molecular functions, and cellular components provide insights into key pathways influenced by DEGs.

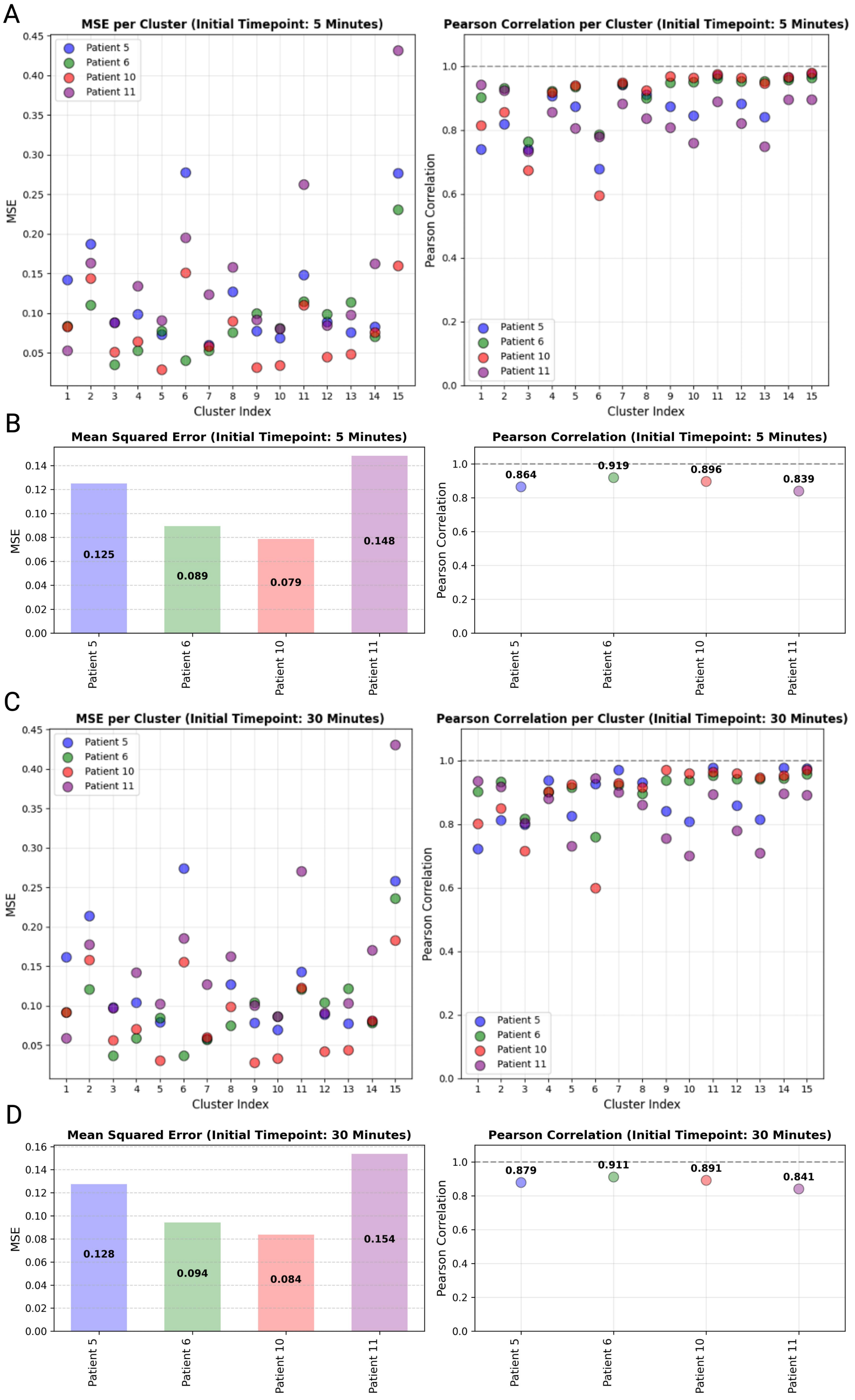


Figure 2: **Patient-specific prediction accuracy.** Comparison of prediction accuracy across individual test patients for the post-surgical initial timepoints of 5 minutes and 30 minutes. Panels A and C show the Mean Squared Error (MSE) and Pearson correlation coefficients between the true and predicted gene expression profiles for each cluster at 5 minutes and 30 minutes, respectively. Panels B and D display the averaged MSE and averaged Pearson correlation across clusters for 5 minutes and 30 minutes, respectively.
